# Force-dependent binding of vinculin to α-catenin regulates cell-cell contacts stability and collective cell behavior

**DOI:** 10.1101/117762

**Authors:** Rima Seddiki, Gautham Hari Narayana Sankara Narayana, Pierre-Olivier Strale, Hayri Emrah Balcioglu, Grégoire Peyret, Mingxi Yao, Anh Phuong Le, Lim Chwee Teck, Jie Yan, Benoit Ladoux, René-Marc Mège

## Abstract

The shaping of a multicellular body and repair of adult tissues require fine-tuning of cell adhesion, cell mechanics and intercellular transmission of mechanical load. *Adherens* junctions (AJs) are the major intercellular junctions by which cells sense and exert mechanical force on each other. However, how AJs adapt to mechanical stress and how this adaptation contributes to cell-cell cohesion and eventually to tissue-scale dynamics and mechanics remains largely unknown. Here, by analyzing the tension-dependent recruitment of vinculin, α-catenin and F-actin as a function of stiffness, as well as the dynamics of GFP-tagged wild-type and mutated α-catenins, altered for their binding capability to vinculin, we demonstrate that the force-dependent binding of vinculin stabilizes α-catenin and is responsible for AJ adaptation to force. Challenging cadherin complexes mechanical coupling with magnetic tweezers, and cell-cell cohesion during collective cell movements, further highlight that tension-dependent adaptation of AJs regulates cell-cell contact dynamics and coordinated collective cell migration. Altogether, these data demonstrate that the force-dependent α-catenin/vinculin interaction, manipulated here by mutagenesis and mechanical control, is a core regulator of AJ mechanics and long-range cell-cell interactions.

**Summary statement:** Combining cell biology and biomechanical analysis, we show here that the coupling between cadherin complexes and actin trough tension-dependent α-catenin/vinculin association is regulating AJ stability and dynamics as well as tissue-scale mechanics.

## Introduction

*Adherens* junctions (AJs) contribute both to tissue stability and dynamic cell movements. The cadherin-catenin adhesion complex is the key component of AJ that bridges neighboring cells and the actin-myosin cytoskeleton, and thereby contributes to mechanical coupling between cells, which drives both cell assembly stability and dynamic cell movements during morphogenetic and tissue repair events (Collins and Nelson, 2015; Guillot and Lecuit, 2013; Mayor and Etienne-Manneville, 2016; Takeichi, 2014). Central to this process is the dynamic link of the complex to actin filaments (F-actin) (Mege and Ishiyama, 2017). Cadherin cytoplasmic tail binds to β-catenin, which in turn binds to the F-actin binding protein α-catenin. α-catenin then links cadherin-β-catenin adhesion complexes to the force-generating actomyosin networks, directly and/or indirectly through other actin binding proteins such as vinculin or afadin.

In addition, mechanotransduction at AJs enables cells to sense, signal, and respond to physical changes in their environment, and the cadherin-catenin complex has emerged as the main route of propagation and sensing of forces within epithelial and non-epithelial tissues (Hoffman and Yap, 2015; Huveneers and de Rooij, 2013; Ladoux et al., 2015; Leckband and Prakasam, 2006). A proposed mechanotransduction pathway involves the myosin II-generated force-dependent change of conformation of α-catenin regulating vinculin recruitment (le Duc et al., 2010; Thomas et al., 2013; Yonemura et al., 2010). At the single molecule force level, it has been shown that α-catenin reversibly unfolds upon forces in the range of those developed by a few myosin motors, allowing the binding of vinculin head (Maki et al., 2016; Yao et al., 2014). The tension-dependent binding of vinculin to α-catenin may thus be central for the adaptation of cadherin-dependent cell-cell contacts experiencing tugging forces in dynamic epithelial layers (Han et al., 2016; Jurado et al., 2016; Kim et al., 2015), and may contribute directly to tissue mechanics and collective cell behavior. However, how this molecular pathway contributes to the dynamics of cell-cell contacts, and allow cells to locally sense, transduce and adapt to environmental mechanical constraint, is not well understood.

Here, we tackled this question by investigating the contribution of the force-regulated interaction between vinculin and α-catenin to AJ dynamics and collective cellular behavior. Indeed, it has been difficult to address this question so-far, at least in mammalian cells, since in addition to the pleiotropic effect of α-catenin loss of function described *in vivo* (Lien et al., 2006; Silvis et al., 2011; Torres et al., 1997; Vasioukhin et al., 2001), the knock-out of the protein in cells *in vitro* lead to the complete inhibition of cadherin-mediated adhesion (Benjamin et al., 2010; Thomas et al., 2013; Vermeulen et al., 1995), as well as to cadherin-independent alteration of actin dynamics, subsequently affecting cell interaction with the ECM (Benjamin et al., 2010; Hansen et al., 2013). To address the role of the tension-dependent association of α-catenin and vinculin, we generated mutant α-catenin proteins either unable to bind vinculin or constitutively bound to vinculin, and analyzed their effect on cell-cell contact stability and collective cell behavior when expressed in α-catenin-depleted epithelial cells.

## Results and discussion

### a-Catenin, vinculin and F-actin recruitment at cell-cell contacts is dependent on intercellular stress in epithelial monolayers

Previous data have shown that vinculin recruitment at cell-cell contacts was dependent on tension. However, these data were obtained either by inhibiting intracellular contractility (le Duc et al., 2010; Yonemura et al., 2010) or by applying external forces (Dufour et al., 2013; le Duc et al., 2010). To directly determine the impact of intercellular stress physiologically generated by cell-born contractile forces, we plated MDCK cells on fibronectin (FN)-patterned polyacrymamide gels of controlled stiffnesses of 4.5, 9 and 35 kPa and looked at the impact on the recruitment of vinculin, α-catenin and F-actin. The junctional recruitment of these three proteins significantly increased with substrate stiffness (**Fig. 1A,B**). This enrichment did not result from increased cellular protein levels as shown by western blot (**Fig. 1C**). Moreover, we did not detect significant changes in junctional accumulation of E-cadherin in function of substrate stiffness’s (1.36 ± 0.12, 1.30 ± 0.11 and 1.33 ± 0.04 A.U. for substrate stiffness’s of 4.5, 9 and 35 kPa, respectively), indicating that there was indeed a specific enrichment in α-catenin, vinculin and F-actin at constant density of junctional E-cadherin. Thus, the junctional recruitments of α-catenin, vinculin and F-actin are positively controlled by the intercellular tension imposed by the matrix stiffness. The staining at cell-cell contacts with an antibody recognizing α-catenin under its open conformation also increased with substratum rigidity, suggesting a central contribution of the tension-dependent conformational change of α-catenin and recruitment of vinculin to the physiological adaptation to force of AJs.

**Figure 1:**
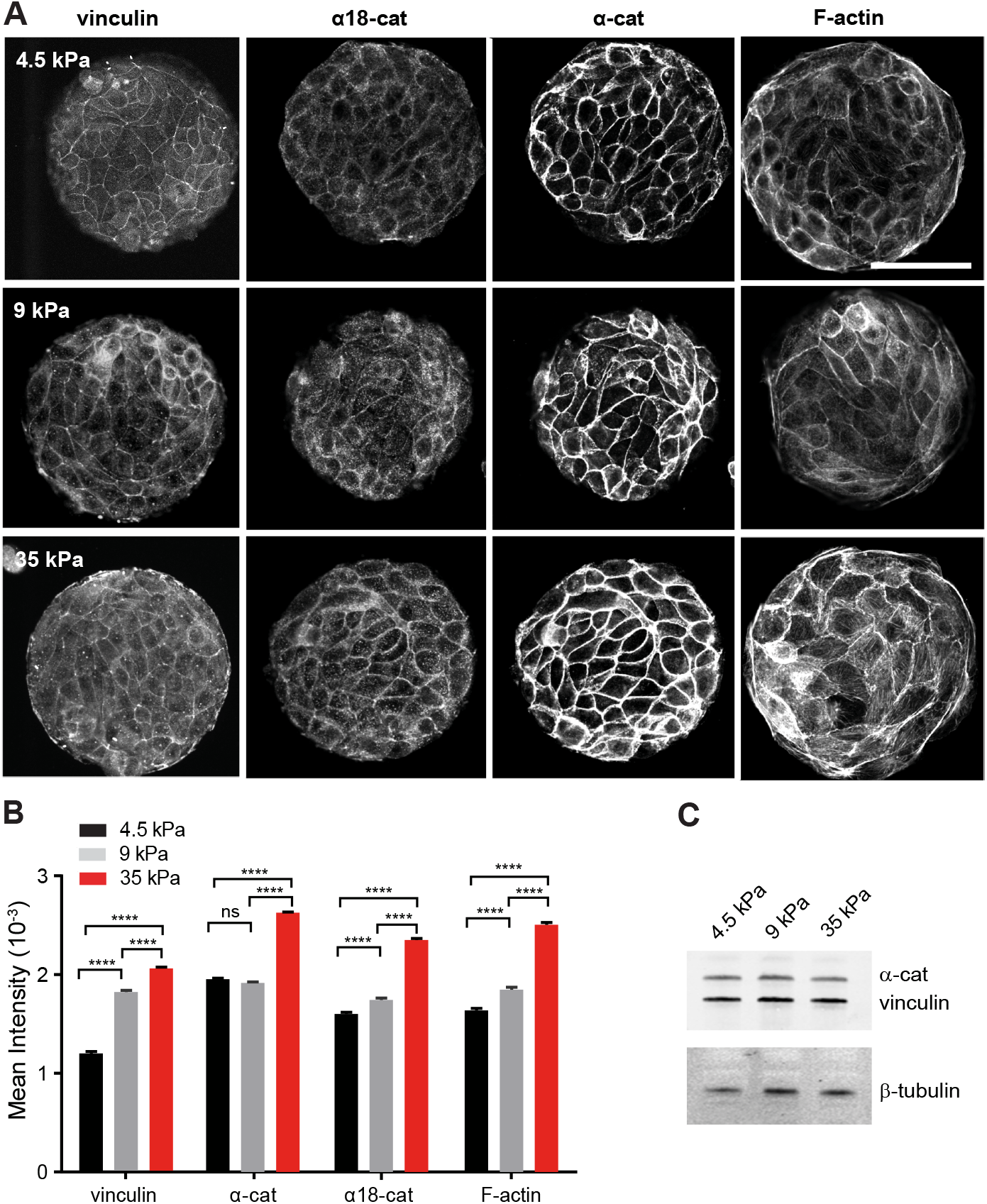
Substrate stiffness-dependent recruitment of α-catenin, vinculin and F-actin at cell-cell contacts. **(A)** MDCK cells were cultured for 24 hours on PAA gels of the indicated stiffness (4.5, 9 or 35 kPa), on which 100 μm diameter disks of FN had been patterned. Preparations were then fixed and immunostained for α-catenin, α18 epitope, vinculin, and F-actin, and then imaged by confocal microscopy (panels show 0.5 μ thick z-projections taken at the level of the apical complexes). Scale bar: 50 μm. **(B)** The histograms represent the mean fluorescence intensities measured for αE-catenin, α18 epitope, vinculin and F-actin stainings as indicated in Materials and Methods (mean ± SEM, n = 640 to 1260 junctions in total per condition, out of 3 independent experiments, **** p< 0.0001, ns: not significant, one-way ANOVA test. **(C)** Western blot analysis of α-catenin and vinculin from protein extracts of cells grown for 24 hours on FN coated PAA gels of 4.5, 9 and 35 kPa rigidity, respectively. α-tubulin was used as a loading control.

### Vinculin binding to a-catenin is not required for the formation of cell-cell junctions *but* stabilizes junctional α-catenin

To address the role of the α-catenin/vinculin interaction in the tension-dependent regulation of cell-cell contacts, we generated α-catenin mutants unable to bind vinculin (α-cat-L344P) or binding constitutively to vinculin (α-cat-Δmod), respectively (**Fig. 2A**). Vinculin binds to α-catenin within the MI domain, and substitution of Lysine 344 by Proline has been reported to impair vinculin binding (Peng et al., 2012; Yao et al., 2014). The MII and MIII domains (residues 509–628) are auto-inhibitory domains interacting with the M1 domain and masking the accessibility of the vinculin binding domain (Desai et al., 2013; Maki et al., 2016). This auto-inhibition is released upon force-dependent stretching, unmasking the vinculin binding domain (Yao et al., 2014). Deletion of residues 509–628 (Δmod mutation) thus generates a α-catenin isoform constitutively binding to vinculin.

**Figure 2:**
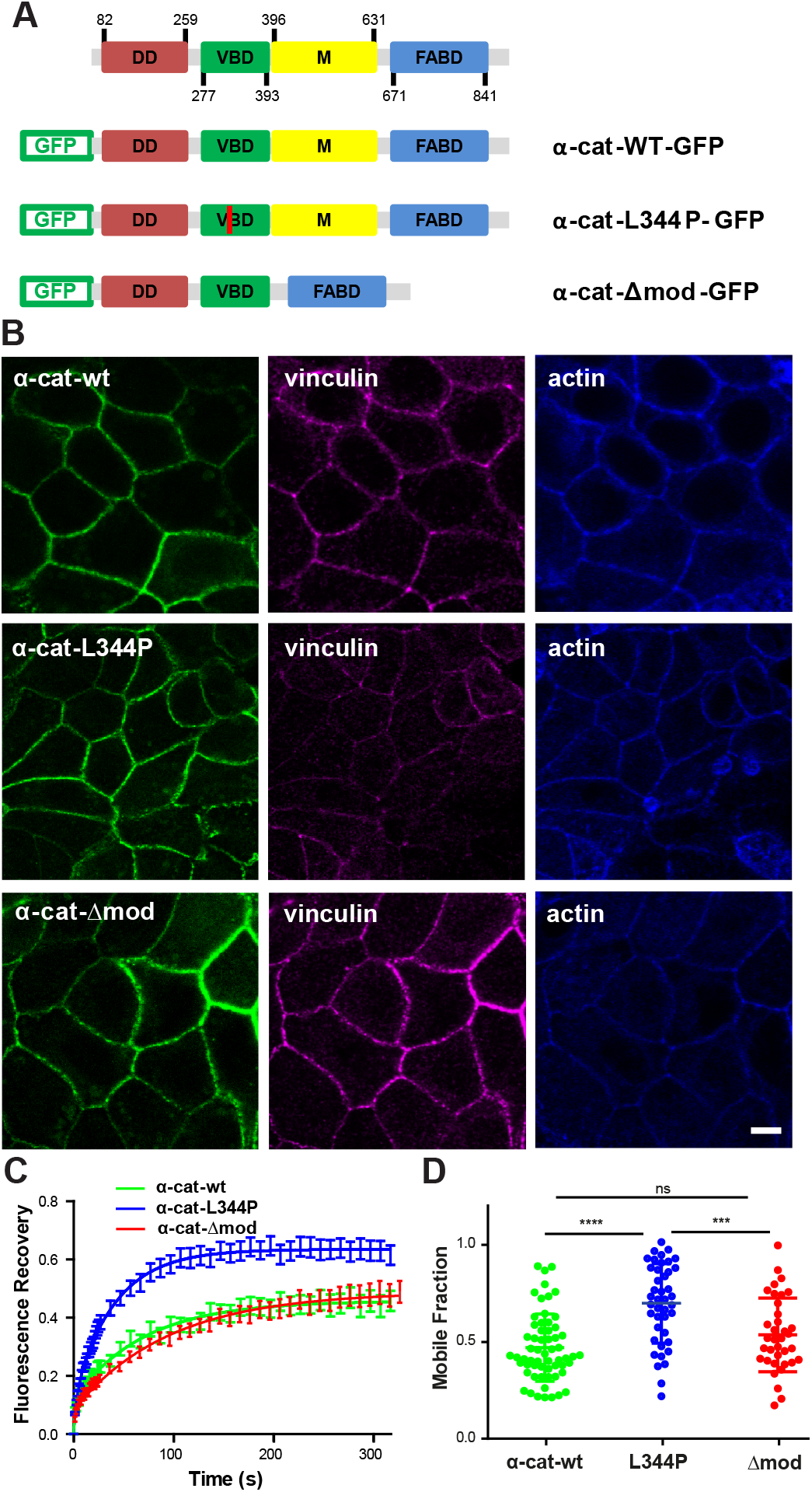
E-cadherin-dependent cell-cell contacts form independently of α-catenin/vinculin interactions. **(A)** Schematics of GFP-tagged wild-type (α-cat-wt-GFP), L344P (α-cat-L334P-GFP) and Δmod (α-cat-Δmod-GFP) αE-catenin constructs. **(B)** Confocal analysis of apical vinculin and F-actin distribution in αE-catenin KD MDCK cells expressing α-cat-wt, α-cat-L334P and α-cat-Δmod grown on glass surfaces. The expression of mutant proteins restored cell-cell contacts, as did the expression of wt α-catenin. Scale bar: 5 μm. **(C)** FRAP experiments were performed on cell-cell contacts of α-cat-wt (green), α-cat-L334P (blue) or α-cat-Δmod (red) expressing cells grown on glass substrates. Mean intensity recoveries over time (± SEM) fitted with a one-term exponential equation (n= 50 regions of interest out of 3 independent experiments for each condition). **(D)** Mobile fractions extracted from the fits of individual recovery curves (scatter dot plot, mean values ± SD). **** p< 0.0001, *** p< 0.001, ns: non-significant, one-way ANOVA test.

We analyzed the consequences of the expression of these variants on cell-cell contact restoration in α-catenin-depleted MDCK cells that do not form AJs (Benjamin et al., 2010). The expression of α-cat-L344P and α-cat-Δmod restored the formation of cellcell contacts that were indistinguishable from those of wild-type α-catenin expressing cells (α-cat-wt) (**Fig. 2B**). The recruitment of vinculin at intercellular junctions in α-cat-Δmod expressing cells (1.04 ± 0.02, n=20) was significantly higher compared to α-cat-wt expressing cells (0.67 ± 0.01, n=31, p value < 0.0001). and significantly lower in α-cat-L344P expressing cells (0.34 ± 0.06, n=24, p value < 0.0001, one-may ANOVA test), while the recruitment of vinculin at the cell-substratum interface was comparable for the three cell types (**Fig. S1**). Thus, the two forms of α-catenin allow the formation of AJs in confluent MDCK monolayers, despite their impaired interaction with vinculin. The residual accumulation of vinculin at cell-cell contacts in α-cat-L344P cells may result from the interaction of vinculin with β-catenin reported previously (Peng et al., 2010; Ray et al., 2013). However, we also we measured a ~30 % residual vinculin staining at cell-cell contacts of α-cat-wt expressing cells after 2 hours of blebbistatin treatment (**Fig. S2**). Thus, the incomplete disappearance of junctional vinculin signal both in blebbistatin-treated α-cat-wt and in α-cat-L344P expressing cells may merely be explained by the local contribution of cytoplasmic vinculin staining. In contrast, blebbistatin treatment had no significant effect on the accumulation of vinculin at cellcell contacts of α-cat-Δmod expressing cells (**Fig. S2**), validating the effect of the Δmod mutation on tension-dependent vinculin binding.

To assess the effect of the alteration of the tension-dependent binding of vinculin to α-catenin on the stability of AJs, we performed FRAP experiments on GFP-α-catenin at cell-cell contacts. The fluorescence recovery was similar for wt and Δmod α-catenin with ~ 50 % the molecules in the fast recovering, diffusion-limited fraction (**Fig. 2C, D**). However, the fluorescence recovery for the L344P mutant was significantly increased with a mobile fraction of α-catenin that was doubled (**Fig. 2C, D**) without changes in half recovery time (**Fig S2A**). Interestingly, E-cadherin stability at cell-cell contacts was barely dependent on the binding of α-catenin to vinculin (**Fig. S3A, B**). α-catenin is however a complex molecule (Kobielak and Fuchs, 2004) and mutations analyzed here may have perturb binding sites for others partners. Thus, to exclude this hypothesis we analyzed the mobility of α-cat-wt (fused to mCherry) in MDCK vinculin knockdown cells (Sumida et al., 2011). We found that α-cat was also strongly destabilized in these conditions with a mobile fraction of 0.72 ± 0.06. Thus, vinculin binding to α-catenin is dispensable for the formation of intercellular contacts *per se*, but is required for the stabilization of junctional α-catenin in confluent monolayers. Vinculin binding may stabilize α-catenin by preventing its re-folding following its force-dependent unfolding (Yao et al., 2014). This may via a force-dependent global conformational switch of α-catenin shift α-catenin and the cadherin-catenin complex into a strongly F-actin bound state, thereby creating a self-reinforcing system for strong linkage of the complex to the actin cytoskeleton (Buckley et al., 2014; Ladoux et al., 2015). Vinculin may also stabilize α-catenin at AJs by providing additional binding interfaces between cadherin complexes and F-actin, thanks to the F-actin binding domain of vinculin (Thomas et al., 2013; Yonemura et al., 2010). Previous studies indeed indicate that the anchoring of cadherin complexes to F-actin by the actin binding site of vinculin in the absence of α-catenin actin binding domain is sufficient to restore E-cadherin-dependent cell contacts (Chen et al., 2015; Jurado et al., 2016; Thomas et al., 2013) and that these junctions are even more stable than wt AJs (Chen et al., 2015). Vinculin incorporation in the cadherin-catenin adhesion complex may thus increase its stability. Whatever is the exact molecular mechanism, vinculin binding under the control of tension-dependent conformational switch of α-catenin may explain the observed substrate rigidity-dependent stabilization of α-catenin, vinculin and F-actin at AJs.

### Binding of vinculin is required for the tension-dependent stabilization of α-catenin at cell-cell contacts

To further determine the requirement of the α-catenin/vinculin interaction in the tension-dependent adaptation of cell-cell contacts, we analyzed the dynamics of junctional α-cat-wt, α-cat-L344P and α-cat-Δmod in cells seeded on PAA substrates of 4.5 or 35 kPa stiffnesses (**Fig. 3**). FRAP experiments performed on α-cat-wt expressing cells revealed a significant stiffness-dependent change in junctional α-catenin dynamics, with a higher mobile fraction on the more compliant substrate (**Fig. 3A, C**). At the opposite, the dynamic of the α-catenin mutants were independent of substrate stiffness (**Fig. 3B, C**). The mobile fraction α-cat-L344P was similar on both substrates and comparable to the mobile fraction value observed for the wt α-catenin on soft substrate. The mobile fraction of junctional α-cat-Δmod was also independent of the substratum compliance, but was significantly lower and comparable to the value obtained for wt α-catenin on stiff substrate. The characteristic recovery half times were not significantly different among these different conditions (**Fig S2B**). The dynamics of E-cadherin did not change significantly with substrate compliance, or with α-catenin mutations (**Fig. S3**), in agreement with the observed vinculin-independent recruitment of E-cadherin around cell-bound E-cadherin-coated beads (le Duc et al., 2010). These data show that the molecular stability of α-catenin at AJs is mechanosensitive, and that this mechanosensitive stabilization requires the binding to vinculin. Altogether, they suggest that the tension-dependent binding of vinculin to α-catenin regulates the stiffness-dependent stabilization of cadherin adhesion complexes at cell-cell contacts.

**Figure 3:**
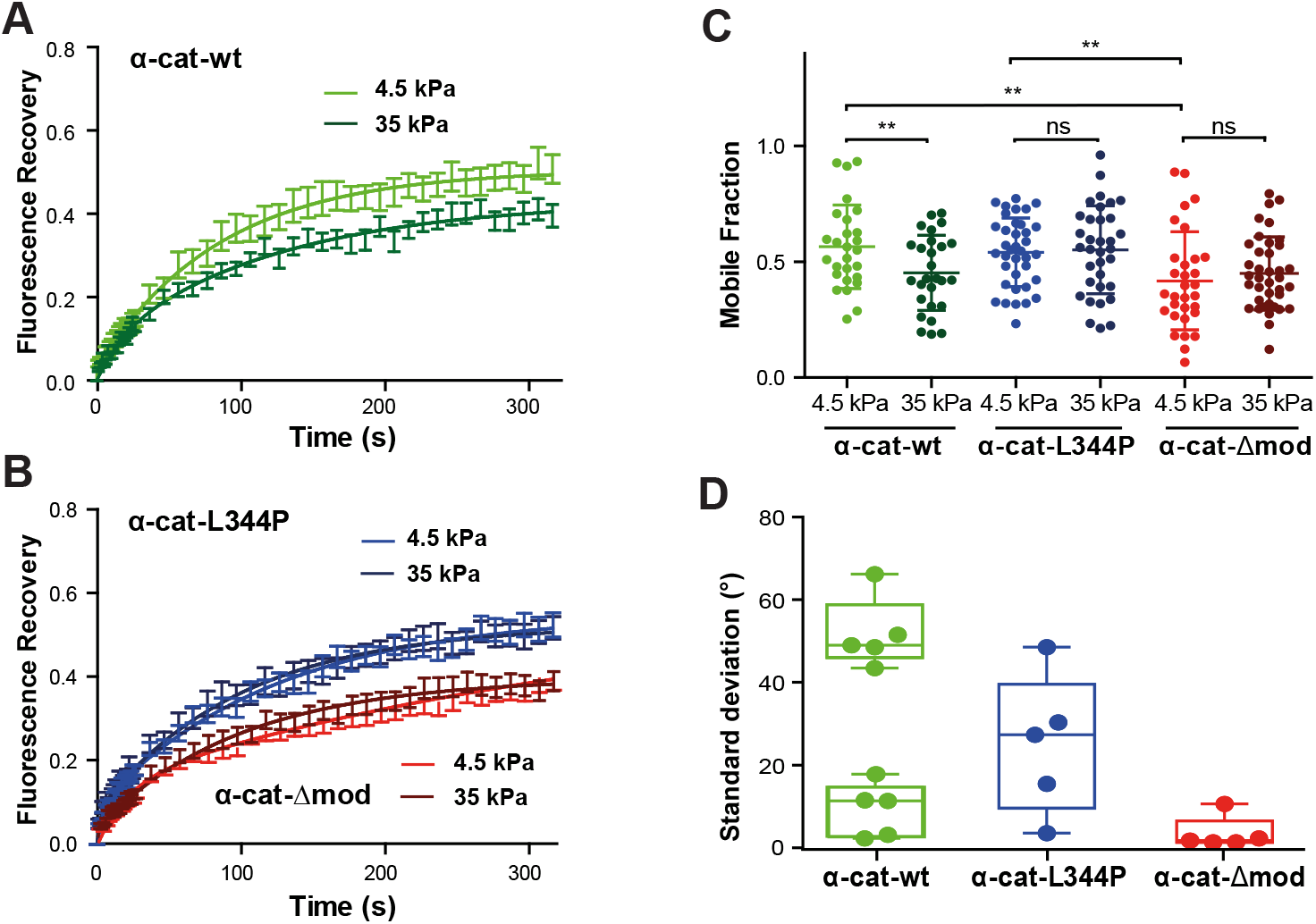
Binding of α-catenin to vinculin is required for its tension-dependent stabilization at cell-cell contacts. **(A, B)** MDCK α-catenin-KD cells expressing GFP-tagged α-cat-wt (green), α-cat-L334P (blue) or α-cat-Δmod (red) were cultured for 24 hours on 4.5 (light colors) or 35 kPa (dark colors) PPA gels before FRAP experiments were performed. Graphs represent mean GFP fluorescence recovery over time (± SEM, n= 50 out of 3 independent experiments for each condition) fitted with a one-term exponential equation. **(C)** Mobile fraction values (scatter dot plot, mean values ± SD) extracted from the fits of individual recovery curves considered in panels **A** and **B**. ** p< 0.01, ns: non-significant, two-way ANOVA test. Notice the non-significant differences in mobile fraction values observed for the mutant proteins on soft and stiff substrates, contrasting with the significant decrease in mobile fraction observed for the wt protein as a function of increasing substrate compliance. **(D)** Magnetocytometry applied on Ecad-Fc coated bead doublets bound to the surface of MDCK cells expressing α-cat-wt, α-cat-L334P and α-cat-Δmod mutants. The histogram reports the mean values of the standard deviation of the bead fluctuation angles.

### Vinculin/α-catenin association controls E-cadherin coupling to cortical actin

To test whether vinculin binding controls the mechanical coupling of cadherin complexes to the underlying actin cytoskeleton, we performed magneto-cytometry experiments using super-paramagnetic Ecad-Fc-coated bead bound on α-cat-wt, α-cat-L344P, α-cat-Δmod expressing cells. Torque were applied to bound beads by rotating a pair of permanent magnets 360 degrees in both clockwise and counter clockwise direction, and the standard deviation of fluctuation angles of the beads were obtained to quantify the response to the applied torque (**Fig. 3D**). This measurement can be considered as a proxy of the stiffness of the mechanical link between cadherins and the cell cortex (le Duc et al., 2010). For cells expressing α-cat-wt, bead fluctuation angles showed two populations; half the beads loosely attached (mean s.d. = 51.8 ± 3.8 °) and the other half strongly coupled (mean s.d. = 9.2 ± 2.9 °). Thus, E-cadherin beads bound to wt α-catenin expressing cells were either tightly or loosely coupled to the cortical cytoskeleton, indicating a complex regulation of the linkage between cadherin complexes and the cell cortex. Whether the two states were associated to differences in the lifetime of the bead-cell contact or to specific sites of binding of the bead on the cell surface are questions that could not be addressed with the present approach. In contrast, the mean s.d. of the bead fluctuation angle was centered to 25.1 ± 7.5 degrees for cells expressing α-cat-L344P, indicating a very loose coupling to the cell cortex. At the opposite the mean s.d. was close to 0 (3.5 ± 1.8 °) for cells expressing α-cat-Δmod, indicating a very stiff link of cadherins to the cortical actin (**Fig. 3D**). These results demonstrate that the binding of α-catenin to vinculin is required for efficient mechanical coupling of cadherin-catenin complexes to the underlying cortical cytoskeleton. Altogether, our findings suggest that the mechanosensitive α-catenin/vinculin interaction, by contributing to tension-dependent cell-cell contact stability, regulate tissue mechanics.

### Vinculin/α-catenin association controls collective cell dynamics

While the role of α-catenin in epithelial tissue mechanics and collective cell behavior is well established, (Bazellieres et al., 2015; Doxzen et al., 2013; Vedula et al., 2012), the contribution of the tension-dependent association of vinculin to α-catenin remains unknown. To address the specific role of the binding of vinculin to α-catenin in epithelial tissue mechanics, we analyzed and compared the dynamics of confluent monolayers of mutant expressing cells, α-cat-wt expressing cells and parental α-cat KD cells monolayers, seeded on 500 μm Ø circular patterns coated with FN (**Fig. 4A, Suppl. Videos 1 to 4**). α-cat-Δmod and α-cat-wt cells displayed slow and coordinated movements. At the opposite, α-cat KD and α-cat-L344P cells behave as mesenchymal-like cells, displayed more rapid and uncoordinated motions as described previously for α-cat KD cells (Vedula et al., 2012). The heat map of velocity fields obtained by Particle Imaging Velocimetry (PIV) analysis further revealed higher velocities for α-cat KD and α-cat-L344P cells than for α-cat-Δmod and α-cat-wt cells (**Fig. 4B**). The average velocities were the highest for α-cat KD cells, then significantly decreased for α-cat-L344P cells, and finally for α-cat-wt and α-cat-Δmod cells (**Fig. 4C**). The spatial velocity correlation which refers to the mean distance at which velocity vectors are oriented in the same direction length was further extracted as a quantitative parameter of cell movement coordination. α-cat KD cells exhibited low coordinated motion with a correlation length of 50 μm as opposed to values around 150–200 μm for wt MDCK cells, as reported by (Vedula et al., 2012). Correlation lengths were comparable for α-cat-Δmod and α-cat-wt cells. However, the correlation length was significantly lower for α-cat-L344P than for α-cat-wt cells denoting an altered coordinated behavior as in the case of α-catenin knock down (**Fig. 4D**). Collective epithelial cell movements are thus strongly dependent on the ability of α-catenin to bind vinculin.

**Figure 4:**
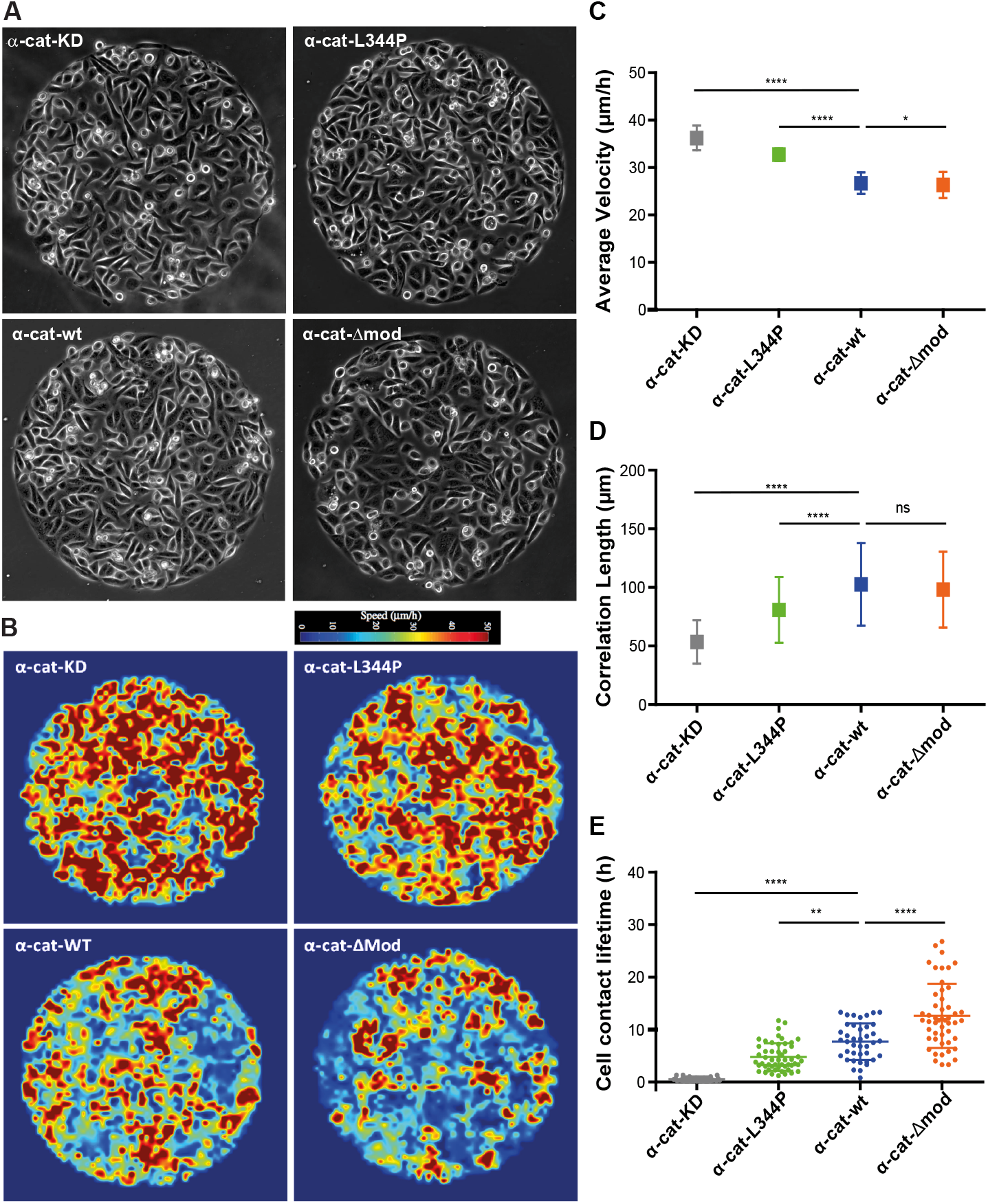
Vinculin/α-catenin association controls collective cell behavior and cell-cell contact lifetime. **(A)** MDCK cells silenced for α-catenin (α-cat KD), as well as cells expressing α-cat-L334P, α-cat-Δmod or wt α-catenin (α-cat-wt), were seeded on 500 μm Ø FN patterns and phase contrast imaged for 24–36 hours (still images of supplementary videos 1 to 4). The collective behavior of cell monolayers was analyzed by PIV over 6 hours providing heat maps of instantaneous local velocities **(B)**. Mean velocities **(C)** and correlation lengths **(D)** characteristic of each cell type were then extracted from these instantaneous velocity map (mean values ± SD) out of 3 independent experiments, n = 360 frames analyzed per condition, derived from 10 patterns per condition coming from 3 independent experiments. **** p<0.0001, * p<0.1, ns: non-significant, one-way ANOVA test. **(E)** Mean lifetime of individual cell-cell contacts measured for each cell type (scatter dot plot: mean ± SD, n = 30 cell doublets for α-cat KD and α-cat-wt, and n= 51 for α-cat-L334P, α-cat-Δmod, out of ≥ 4 patterns derived from ≥ 2 independent experiments for each condition, **** p<0.0001, ** p<0.01, one-way ANOVA test).

An increased stability of cell-cell contacts, by limiting neighbor exchange within the monolayer, may directly explain a more coordinated cell behavior and vice versa. This was confirmed by quantifying the effect of vinculin-binding capability of α-catenin on the lifetime of cell-cell contacts in monolayers coexpressing RFP-Ftractin (**Fig. 4E, Suppl. Videos 5 to 8**). While cell-cell contact lifetime was of less than 1 hour for α-cat KD cells, it was of 4.7 ± 2.6 h for α-cat-L344P cells and increased to 7.7 ± 3.4 h and 12.6 ± 6.1 h for α-cat-wt and α-cat-Δmod cells, demonstrating that the binding of vinculin to α-catenin is required for maintaining stable cell-cell contacts during collective motion of epithelial cells. To support further that the α-catenin/vinculin interaction is responsible for cell-cell contact stability, we analyzed cell-cell contact lifetime in vinculin KD MDCK monolayers (Sumida et al., 2011). Vinculin KD MDCK cells had very short cellcell contact lifetime with a mean value of 2.7 ± 1.7 h in between the one measured for α-cat KD cells and the one measure for α-cat L344P cells. However, due to the strong effect of vinculin knockdown on cell-substratum adhesion, it was not possible to draw conclusion on the effect on correlation length and velocity of the monolayer movement. Thus, we show here that coordinated motion of epithelial cells is dependent on the ability of α-catenin to bind vinculin, which drastically increase cell-cell contact lifetime and correlated movements.

Altogether our results show that the force-dependent interaction between α-catenin and vinculin is crucial for epithelial cells to develop stable but adaptive cell-cell contacts in response to the mechanical resistance of their environment as well as for long-range cell-cell interactions and tissue-scale mechanics. The fact that the association between the two proteins, manipulated here by mutagenesis and mechanical control, has a direct incidence on collective cell movements put the core α-catenin and vinculin mechanosensing machinery at the center of the control of morphogenetic processes.

## Material and Methods

### Cell culture

Madin-Darby canine kidney (MDCK, from ATCC), MDCK αE-catenin knockdown (Benjamin et al., 2010) and MDCK vinculin knockdown cells (Sumida et al., 2011) were maintained in DMEM/Glutamax 10% fetal bovine serum (FBS), 100 μg/mL penicillin/streptomycin at 37°C in 5% CO_2_. Cells were electroporated using Amaxa system (Nucleofector Device, Lonza) under the following conditions 24 hours prior to experiments: 1.10^6^ cells, kit L, 5 μg of DNA.

### Expression vectors

The wt GFP-α-catenin expression vector coding for GFP fused in N-terminal of mouse αE-catenin was described previously (Thomas et al., 2013). L344P GFP-α-catenin wt mCherry-α-catenin was derived from the wt construct by PCR and DNA ligation. Δmod GFP-α-catenin was obtained by deleting the sequence coding for residues 509–628 (domains MII and MIII according to (Desai et al., 2013)).

### Immunofluorescent staining

Cells were fixed for 15 min with PBS, 4% formaldehyde, permeabilized with 0.15% Triton X-100 in PBS for 5 min, then incubated with primary antibodies (anti-vinculin mouse monoclonal antibody (1/200, clone 7F9, Millipore), rabbit polyclonal anti-α-catenin and anti-β-catenin (1/400, Sigma), rat α18 monoclonal anti-αE-catenin antibody (1/200, A. Nagafuchi, Kumamoto Univ.) in PBS, 1.5% BSA for 2 hours at room temperature. The secondary antibodies (Jackson Laboratories) and Alexa-643 phalloidin (Molecular Probes) were incubated for 1 hour in PBS-BSA.

### Polyacrylamide substrates

The preparation of PAA was adapted from (Pelham and Wang, 1997). The PAA gel formulations were prepared from solution of 2% bis-acrylamide and 40% acrylamide (Bio-RAD) in 10 mM HEPES. Bis-acrylamide and acrylamide were first combined and allowed to rest for 15 min. Next, 1/1000 total volume TEMED (Bio-RAD) and 1/100 total volume 10 % Ammonium Persulfate (Bio-RAD) were added to the PAA solution together with 2 mg/ml NHS-ester (Sigma) to allows the formation of covalent bonds between PAA and FN so that FN remains attached to the PAA gel after removal of the stamped coverslip. The concentrations of acrylamide and bis-acrylamide were adjusted to obtain Young’s moduli of 4.5, 9 and 35 kPa, according to (Tse and Engler, 2010).

### Micropatterning

PDMS (Poly-dimethylsiloxane) stamps were molded and cured as described previously (Vedula et al., 2014) on silicon wafers (IEMN (Lille, France) or MBI (NUS, Singapore)). A thin layer of PDMS was spin coated over a 35 mm plastic petri dish and cured at 80°C for 2 hours, then exposed to UV for 15 min in a UVO cleaner (Jelight Company Inc.). PDMS stamps were incubated with a solution of FN/Cy3-conjugated FN (10/1 ratio, 50 μg/ml, Millipore) for 45 min, washed with water and air dried, then gently pressed against the surface to transfer FN. Regions outside the patterns were blocked with 0.2% Pluronics F-127 (Sigma) for 1 hour and washed 3 times with PBS.

Alternatively, polyacrylamide (PAA) substrates were patterned. For this, air dried FN-coated PDMS stamps were brought into contact with a cleaned 22 mm by 22 mm glass coverslip. A second cleaned coverslip, serving as the base for the gel, was silanized by dipping in ethanol solution containing 2% (v/v) 3-(trimethoxysilyl) propyl methacrylate silane (Sigma) and 1% (v/v) acetic acid for 5 min. 20 μl of PAA solution were placed between the silanized coverslip and the FN-stamped one. After 45min of polymerization the patterned coverslip was removed and the coverslip with the gel, on which patterned FN had been transferred, placed in a solution of 10 mM HEPES.

### Videomicroscopy

Transfected cells were seeded at subconfluency on 500 μm Ø FN patterns in DMEM medium, 10 % FCS, allowed to attach 4 hours, washed with culture medium, then live imaged (phase contrast and fluorescence) at low magnification (10x, Biostation^™^, Nikon) every 10 min for 36 hours. The monolayer behavior was analyzed from the time cells occupy entirely the patterns by PIV (Particle Imaging Velocimetry) performed on phase contrast images, to obtain instantaneous velocity fields and correlation length in motion within the cellular disks (Strale et al., 2015).

### Florescence image acquisition and analysis

Preparations were imaged either with a wide field fluorescent microscope (Olympus IX81) equipped of 60x oil immersion objective and a Coolsnap HQ CCD camera, or with a Leica Sp5-II confocal microscope with a 63 X oil immersion objective with a distance of 0.5 μm between each plane in the z-stacks. Protein recruitments at cell-cell contacts were determined on z-stack confocal images encompassing the apical cell domain of the monolayer (3 most apical stacks) analyzed the “surfaces” module of Imaris software, applying background subtraction, thresholding of junction area and removal objects outside junctions. Then the mean intensities of the different staining were measured in the volume of junctions.

### Fluorescence Recovery after Photobleaching

Fluorescence recoveries after photobleaching were performed at 37°C 24 hours posttransfection, using the Leica Sp5-II setup. Argon laser power was set at 50 % for; AOTF (488) output power was set at 10 % for pre and post-bleach acquisition and at 100 % for bleaching. Fluorescence was acquired for 6.5 s at a frequency of 0.8 s^−1^, over a whole field, then a Ø 2 μm ROI was bleached for 3 s, followed by acquisition over the whole field for 26 s at a frequency of 0.8 s^−1^, then for 300 additional seconds at a frequency of 0.1 s^−1^. After correction for photobleaching, the normalized recovery of fluorescence was expressed as a ratio of prebleach fluorescence, as reported previously (Lambert et al., 2007). Fluorescence recoveries in function of time were fitted with one-term exponential equation, allowing to extract a plateau value representing the fraction of diffusion-limited molecules (mobile fraction) and a recovery half-time (t_1/2_) proportional to the apparent diffusion coefficient of diffusion-limited molecules (Thoumine et al., 2006). The mobile fraction and the t_1/2_ were determined by fitting the normalized recovery curves using one-phase decay non-linear regression function of the GraphPad Prism 5.01 software.

### Magnetic tweezers cytometry

Ecad-Fc-coated bead doublets attached to the surface of transfected cells were submitted to magnetic twisting using an in-house built magnetic tweezers. Bead doublets were used instead of individual beads to increase the applied torque and facilitate subsequent image analysis. The bead doublets were made by mixing antibody immobilized Ecad-Fc-coated beads with Protein A coated beads therefore only one bead of the doublet binds to the cell through E-cadherin interaction. Bead doublet rotation was captured while the rotation of the magnet was simultaneously recorded. The fluctuation angle of the bead doublet relative to its original direction was calculated to obtain its response to magnet rotation, and the standard deviation of the fluctuation angle was extracted. More specifically, protein A coated paramagnetic beads (2.8 μm from Dynabeads) were washed 3 times and incubated overnight at room temperature with anti-human Fc fragment antibody in 0.1M borate buffer pH 8. After 3 washes, the beads were resuspended in Ca^++^ and Mg^++^ containing PBS and incubated for 3 hours at room temperature with recombinant Ecad-Fc (R&D Systems). These Ecad-Fc coated beads were mixed with equal amount of uncoated protein A beads and incubated for one hour and room temperature to form bead doublets. The bead solution was then resuspended in PBS, 1% BSA, sonicated, and incubated for 20–30 min at 37C, 5% CO_2_ with adherent transfected cells. GFP-positive cells displaying a bead doublet were selected for the magnetic twisting experiment. The torque was applied thanks to a home-made magnetic twisting cytometer (360 degree in clockwise and anti-clockwise direction at a constant speed of 5 degrees / sec). Bead displacement was captured with an inverted microscope equipped with a 50X long working distance objective lens and a camera (Pike from Allied vision) with 100 Hz sampling frequency while the rotation of the magnet was recorded directly. The image sequence of the bead doublets was loaded into imageJ. The image stack was thresholded to only highlight the contours of the bead doublet and other features of the image like cell contours were removed manually. The ‘Directionality’ plugin of the Fiji package was used to calculate the principle orientations of the bead doublet. The fluctuation angle of the bead was matched to the direction of the magnet to obtain its response to magnet rotation. The standard deviation (std) of the fluctuation angle was used as a measure of the bead-cell interaction stiffness. For a bead that follows the magnet directly, denoting a very soft interaction between the bead and the cell cortex, the standard deviation is around 60 degrees; for very strong mechanical coupling between the bead and the cortex this value is close to 0 degrees.

### PIV analysis

PIV analysis of monolayer movement was performed as described in (Petitjean et al., 2010) using MatPIV v. 1.6.1 package and implemented in MATLAB MatPIV – the PIV toolbox for MATLAB-available at http://folk.uio.no/jks/matpiv/Download/index2.html). Cross correlation techniques were performed to compute the displacement vectors at each sub-window by finding their best match at the successive time frame. The analysis was done with 32 × 32 pixels (19 × 19 μm) interrogation windows with an overlap of 50%. The correlation length was then calculated using the formula previously described (Petitjean et al., 2010). Velocity values obtained were the mean of the norm of all velocity vectors extracted from each image. For this study, we consider the mean velocities over a time interval of 6 hours to minimize effects of cell division, time zero being the moment at which cells reach confluency.

### Life time of cell-cell contacts

The appearance and disappearance of individual cell-cell contacts were tracked over time on time-laps of RFP-Ftractin fluorescence movies. The *t* zero was set when two cells initiate a contact and the separation time, the time at which the two cells separated within the monolayer.

### Statistical analysis and curve fitting and image processing

Statistical analysis and curves fitting were performed with GraphPad Prism 5.0 software. Image processing were done in Image J (or Matlab when indicated), then with Photoshop and Illustrator.

## Acknowledgements

This work was supported by grants from CNRS, Université Paris-Diderot (RMM,BL), NUS-SPC joined program (RMM), Fondation ARC (RMM), Human Frontier Science Program (HFSP) grant RPG0040/2012 (B.L., R.M.M.), Agence Nationale de la Recherche (ANR 2010 Blan1515). BL is supported by the European Research Council under the European Union’s Seventh Framework Programme (FP7/2007–2013/ERC grant agreement no. 617233) and the Mechanobiology Institute. R.S. has been supported by a C’nano program Région Ile de France doctoral fellowship and FRM (FDT20140930851). The authors acknowledge the IJM Imagoseine Imaging Facility, member of the France BioImaging infrastructure supported by the French National Research Agency (ANR-10-INSB-04, « Investments of the future »).

